# Simulating the impact of sensorimotor deficits on reaching performance

**DOI:** 10.1101/139857

**Authors:** Sean M. Sketch, Cole S. Simpson, Frédéric Crevecoeur, Allison M. Okamura

## Abstract

The healthy human nervous system accurately and robustly controls movements despite nonlinear dynamics, noise, and delays. After a stroke, motor ability frequently becomes impaired. To provide insight into the relative impact of specific sensorimotor deficits on motor performance, we modeled neural control of reaching with the human upper limb as a near-optimally feedback-controlled two-degree-of-freedom system with biologically based parameters. We added three sensorimotor impairments commonly associated with post-stroke hemiparesis—abnormal joint coupling, increased noise on internally modeled dynamics, and muscular weakness— and examined the impact on reaching performance. We found that abnormal joint coupling unknown to the system’s internal model caused systematic perturbations to trajectories, longer reach durations, and target overshoot. Increasing internal model noise and muscular weakness had little impact on motor performance unless model noise was increased by several orders of magnitude. Many reaches performed by our perturbed models replicate features commonly observed in reaches by hemiparetic stroke survivors. The sensitivity to unmodeled abnormal joint coupling agrees with experimental findings that abnormal coupling (possibly related to internal model errors) is the main cause of post-stroke motor impairment.

## I. Introduction

The healthy human nervous system quickly, accurately, and robustly controls movement in the face of myriad difficulties including nonlinearities, noise, and delays [1]. Despite this adeptness, roughly 5 million individuals become permanently disabled by stroke each year [2]. Researchers and clinicians note that disabled stroke survivors’ nervous systems must contend with additional sensorimotor deficits such as abnormally coupled joints [3]–[5], increased noise in sensing and estimation [6], muscular weakness [7], and spasticity [8]. These difficulties generally correlate with motor impairment, working in concert to degrade motor performance.

The complexity of the neuromuscular system makes it challenging to determine causal relationships between the sensorimotor deficits facing stroke survivors and their resultant motor impairments [9]–[11]. Many techniques have been developed to study how the nervous system controls movements. Imaging techniques (e.g., fMRI), stimulation techniques (e.g., transcranial magnetic stimulation (TMS), deep-brain stimulation, optogenetics), and movement studies (e.g., center-out reaching tasks, perturbed locomotion studies [12], simple games [13]) have all helped us understand the functioning of healthy and impaired nervous systems. Studies using these techniques correlate inputs (e.g., displayed images in fMRI studies, electromagnetic pulses in TMS, mechanical perturbations in movement studies) with neurophysiological outputs (e.g., blood-oxygen contrast, electrical signals in the muscles, bodily kinematics/dynamics). However, motor impairments can stem from sensorimotor deficits at any stage of the control process—sensory encoding, motor planning, or motor execution. These details are difficult to identify using the input-output approach.

Computational models offer the ability to ethically and noninvasively alter the system being studied in a controlled manner (e.g., weakening muscles [14], shifting tendon-attachment points [15]). Such models are useful for testing scientific theories (e.g., central pattern generators [16]), planning clinical interventions (e.g., tendon-transfer surgery [15]), and even designing engineered systems (e.g., McMahon’s tuned track [17]). Some studies have used optimal control—thought to be a good generative model of the healthy human nervous system [1], [18]–[20]—to reproduce human behaviors [18], [21]–[23]. While such models have been used extensively to study the healthy nervous system, they have rarely been applied to understand motor impairment, particularly that of neural origin.

To examine how sensorimotor deficits may impact motor performance, we model the healthy human neuromuscular system as a planar two-degree-of-freedom (DOF) arm controlled by a near-optimal feedback controller with biologically based parameters (e.g., mass, inertia, sensory noise, actuator limits). To explore the behaviors captured by this model, we add three sensorimotor impairments commonly associated with post-stroke hemiparesis: abnormally coupled joints, increased noise on internally modeled dynamics, and muscular weakness. We demonstrate the sensitivity of our model to each of these deficits and show general consistency with experimental assessments of post-stroke hemiparesis.

## II. Methods

We model the healthy neuromusculoskeletal system as a near-optimally feedback-controlled planar arm, summarized in Fig. 1 with variables and parameters listed (including references for specific values) after Section III. We then perturb the model in ways consistent with sensorimotor deficits associated with post-stroke hemiparesis.

**Fig. 1.**
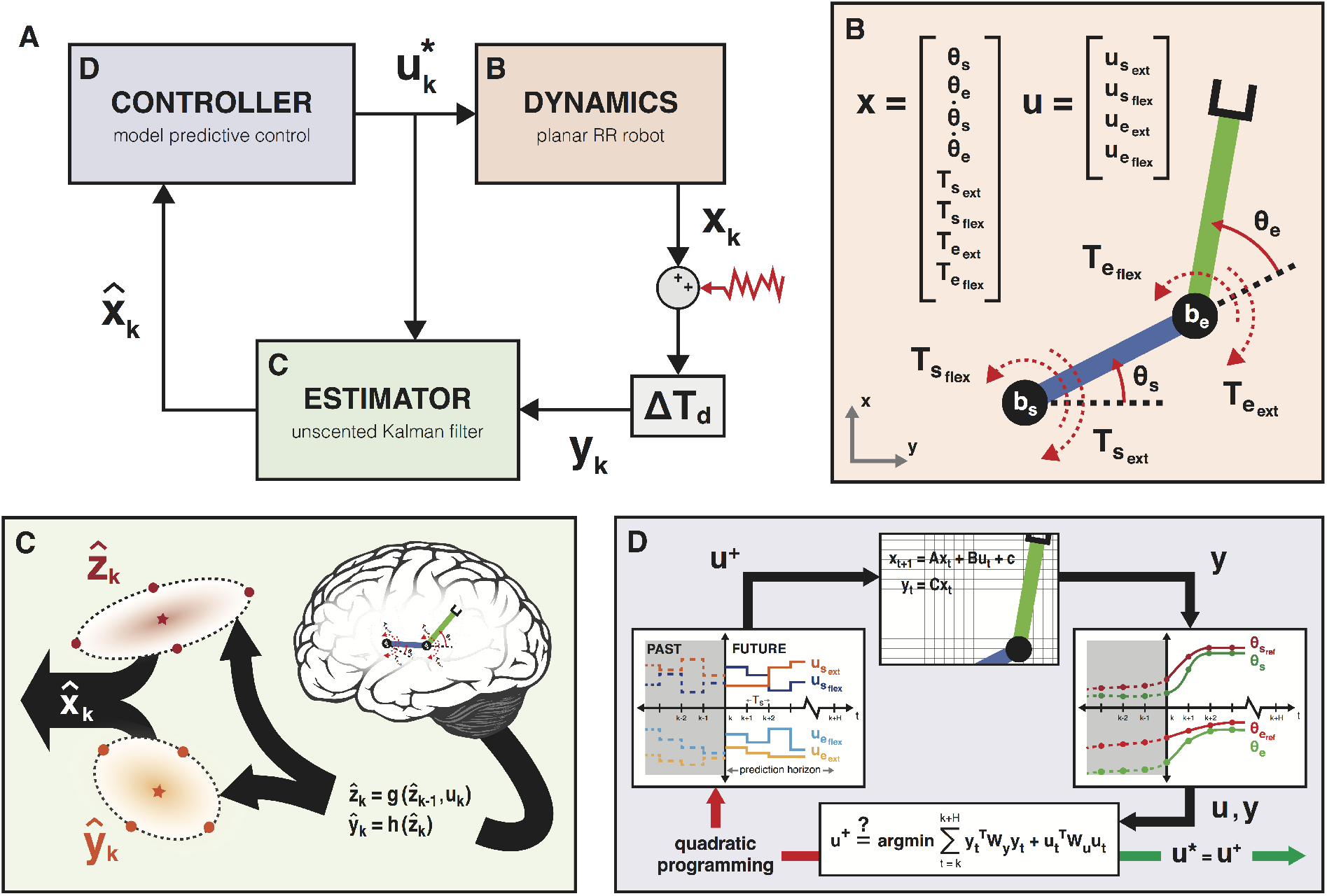
Model overview. We model the neuromuscular system for the arm as a near-optimally feedback-controlled revolute-revolute (RR) robot with biologically based parameters. **Panel A** is a block diagram of the closed-loop system. In addition to the dynamics, estimator, and controller, the model includes a low-pass filter to reflect muscle activation dynamics (not illustrated), sensory noise and bias (jagged red arrow), and time delays thought to represent nerve conduction dynamics and neural computation time (gray block preceding estimator). **Panel B** shows the RR torque-controlled robot and relevant states (joint angles/velocities/torques), inputs (commanded joint torques), and sign conventions. **Panel C** illustrates the unscented Kalman filter used to optimally estimate the state of the arm. The filter combines sensory feedback with predictions made by the brain’s internal model based on the reliability of that information [24]. **Panel D** illustrates the model predictive controller (MPC) used to compute joint torques for the arm. The MPC uses a model of the arm to estimate the effect of candidate torque trajectories and selects the ones that minimize the given cost function over a finite time horizon.

### A. Model

#### 1) Dynamics

We model the arm using the dynamics of a revolute-revolute (RR) robot constrained to move in the plane of the shoulder [18], [20], [25], [26]. The nonlinear differential equation of motion is as follows:

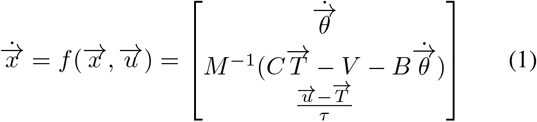

where state 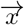 contains the arm’s realized joint angles, velocities, and torques and input 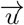 contains commanded joint torques:

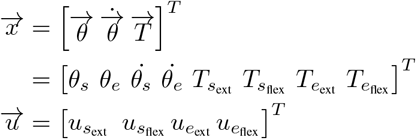

Joint torques are separated based on contribution to flexion (positive) and extension (negative), thereby allowing for cocontractions. In Eqn. 1, *M*, *C*, *V*, and *B* represent the arm’s mass matrix, dynamic coupling matrix, centrifugal and Coriolis effects, and viscous damping (parameters taken from [27] and [28]). *M* and *V* vary nonlinearly with arm state, while *C* and *B* remain constant. To model first-order muscle activation dynamics, we place a low-pass filter with time constant *τ* = 60 ms between the commanded and realized joint torques [28]. Panel B of Fig. 1 defines joint origins and sign conventions for the model; conventions for commanded joint torques (not illustrated) mirror those for realized joint torques.

We numerically integrate the equation of motion over each simulation time step (10 ms) using a variable-time-step medium-order Runge-Kutta method. This gives the following discrete-time state-update equation:

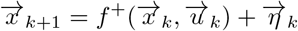

where 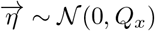 captures white Gaussian process noise with covariance

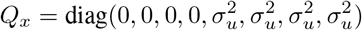

where *σ_u_* = 0.02 N-m is the standard deviation of motor noise [29].

Time delays (estimated to be 60 ms for transcortical pathways) result from nerve conduction dynamics and neural computation. To capture this in our model, we augment the state vector with previous time steps up to the first one observable by the controller [28], [29]. For a delay of *T_d_* time steps, the augmented state 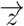 at time step *k* is

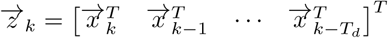

and the corresponding state-update equation is

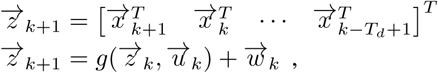

where 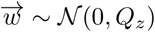 with covariance

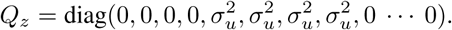

The system senses a noisy and biased version of the delayed arm state 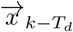 at time step *k*. We model uncertainty in the sensory feedback 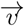 as additive white Gaussian noise and sensory bias 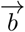 as a linear function of the current arm configuration [30]. This gives the following feedback equation:

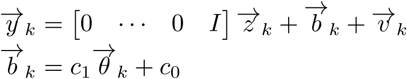

where constants *c*_0_ and *c*_1_ and 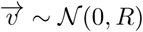 with covariance 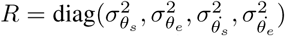 are fit to data from Cusmano *et al*. [6]. Because bias is a function of current state, we can condense the feedback equation as follows:

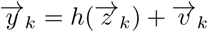

#### 2) Estimation

For control, the neuromuscular system needs reliable information about the arm’s current state. However, at any given time, the system has only delayed and noisy sensory information coming in from the periphery. Just as the (healthy) neuromuscular system appears to optimally control movements, it also appears to optimally estimate the state of the limb [31]. We approximate this optimal estimation using a Kalman filter, merging sensory feedback with state predictions generated by the brain’s internal model:

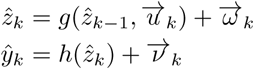

where 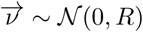 and 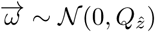 with covariance

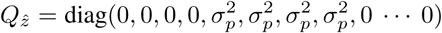

*σ_p_*, the standard deviation of noise on internally modeled dynamics (i.e., internal process noise), is all that differentiates the internal mental model of the arm from the physical plant. We will refer to *σ_p_* as estimation noise.

By accounting for time delay via state augmentation, the filter estimates all states within the interval of the time delay. Given the arm’s nonlinear dynamics, we use an unscented Kalman filter (UKF) [32]. As depicted in panel C of Fig. 1, the UKF uses sampling points to capture the mean and covariance of the predicted state. Unlike the extended Kalman filter, which relies on a linear approximation of the dynamics, the UKF’s sampling points can be exactly propagated through the nonlinear equation of motion. To the best of our knowledge, use of the UKF in computational biomechanics has not been seen in the literature.

#### 3) Control

We use a model predictive controller (MPC) to determine near-optimal joint torques for the arm [33]. This MPC uses the brain’s internal model of the arm to compute a dynamically consistent, optimal torque trajectory over a finite time horizon while accounting for limits on joint positions and torques (from Chackwick *et al*. [34] and van Dijk *et al*. [35]). Previous work on human motor control suggests that movements generally balance error minimization with effort minimization [36]. This is reflected in our finite-time-horizon optimization:

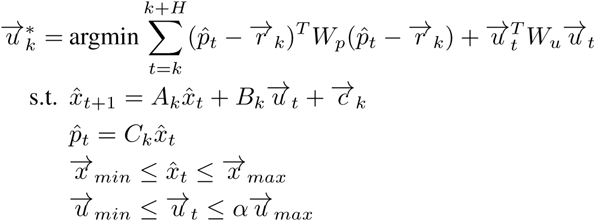

In the cost function, *H* is the number of time steps in the finite horizon, 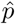 is the internal model’s hand position, 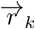 is the target location specified in Cartesian coordinates (constant during the reaches performed here), *W_p_* and *W_u_* are weighting matrices, and 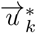 is the resultant optimal torque trajectory. We solve the MPC optimization problem using the Multi-Parametric Toolbox (MPT3 [37]), which requires linear or affine equality constraints. Therefore, the first constraint equation is a linearization (in this case, affinization) of the internal model’s dynamics 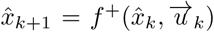 and the output equation 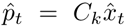 is a linearization of the internal model’s forward kinematics. Both linearizations are first-order Taylor-series approximations about the current estimated arm state and known commanded joint torques, which is only appropriate for *t* “close to” *k*; we assume that this holds over the finite horizon. By linearizing the forward kinematics, we track a reference in Cartesian instead of joint space. Finally, the parameter *α* in the inequality constraint for control effort is a scalar between 0 and 1; it provides a rough approximation of arm strength. We re-plan joint torques every 30 ms.

### B. Model Perturbations

We perturbed the healthy model in ways that align with sensorimotor deficits commonly observed following stroke.

#### 1) Abnormal joint coupling

Chronic stroke survivors frequently exhibit abnormal coupling between joints [3], [11], [38]. To examine the effects of this coupling on motor performance, we changed the coupling matrix *C* in the arm’s equations of motion (Eqn. 1) from an identity matrix to one with off-diagonal elements extracted from data published by Dewald *et al*. [3]. As alluded to in Sections II-A.2 and II-A.3, our simulations use two arm models: a “physical” arm as the plant and the brain’s internal arm model for estimation and control. Though stroke primarily affects the brain, previous studies have shown substantial improvements in upper-extremity motor performance by adding mechanical assistance to the arm [11]. This improvement suggests that cortical planning and estimation remains intact while the issues stem from sub-cortical signal transmission and the execution of motor commands. Because coupling the joints in both the plant and internal model does not result in impaired reaches (tested, but not shown), we explored the effect of joint coupling in the plant only. We simulated center-out reaches to 8 different targets with both a “healthy” (unperturbed) plant and one with abnormal joint coupling.

#### 2) Increased estimation noise

In addition to motor deficits, stroke survivors have difficulty estimating the state of their limbs [6]. Although Cusmano *et al*. [6] demonstrated a stroke-induced increase in proprioceptive bias and noise, simulations with this deficit alone do not display any impairment (tested, but not shown). Poor estimation is more likely caused by increases in the internal model’s prediction error. This is achieved by increasing estimation noise (i.e., process noise within the Kalman filter). For healthy reaches, we assume *σ_p_* = 0.02 N-m [29]. To examine the effect of poor estimation, we simulated reaches to targets with *σ_p_* = 0.2, 2, and 8 N-m.

#### 3) Muscular weakness

Hemiparetic subjects are commonly noted to have muscular weakness. We modeled the effect of this weakness by reducing the maximum torque constraint in the optimization problem to 80% (*α* = 0.8), 40% (*α* = 0.4), and 10% (*α* = 0.1) of the healthy values. We simulated reaches to targets for each weakness level.

## III. Results and Discussion

Planar reaches simulated with our model exhibited characteristics of healthy human motor control: straight reaching trajectories with bell-shaped velocity profiles (Figs. 2 and 3, blue lines). When we systematically perturbed the model in ways associated with post-stroke sensorimotor deficits, each of the perturbations produced changes in motor performance subjectively characteristic of hemiparetic reaches. Although we could have perturbed the model in a subtractive (removing deficits from a fully deficient model) as opposed to additive (introducing deficits to the healthy model), it is unclear how to define “fully deficient.” Moreover, at this stage of model development—prior to rigorous validation— the additive model is at least as instructive as the subtractive one. Subtractive modeling will be important to consider when validating the model, as well as when using the model to assist with the design of rehabilitative devices.

Our simulations indicate that abnormally coupled joints cause systematic direction-dependent perturbations to straight reaching trajectories, target overshoot, and longer settling times. These trajectories are subjectively similar to reaches performed by hemiparetic subjects [39], [40]. Moreover, the sensitivity of the model to the coupling agrees with experimental studies suggesting that abnormal joint or muscle coupling causes the bulk of motor impairments after stroke [10], [11], [40]. The simulation results presented in Fig. 2 show the effects of coupling efferent motor commands without allowing the brain to internally account for this coupling (i.e., capture this coupling in its internal model). For this reason, such a perturbation might be most representative of an acute stroke patient, for whom the mapping between neural commands and muscles has been altered, but the nervous system has not yet had time to learn the new mapping. This type of perturbation might also be representative of stroke patients who were unable to learn an altered mapping between neural commands and muscles during the critical recovery period following the stroke event [41].

**Fig. 2.**
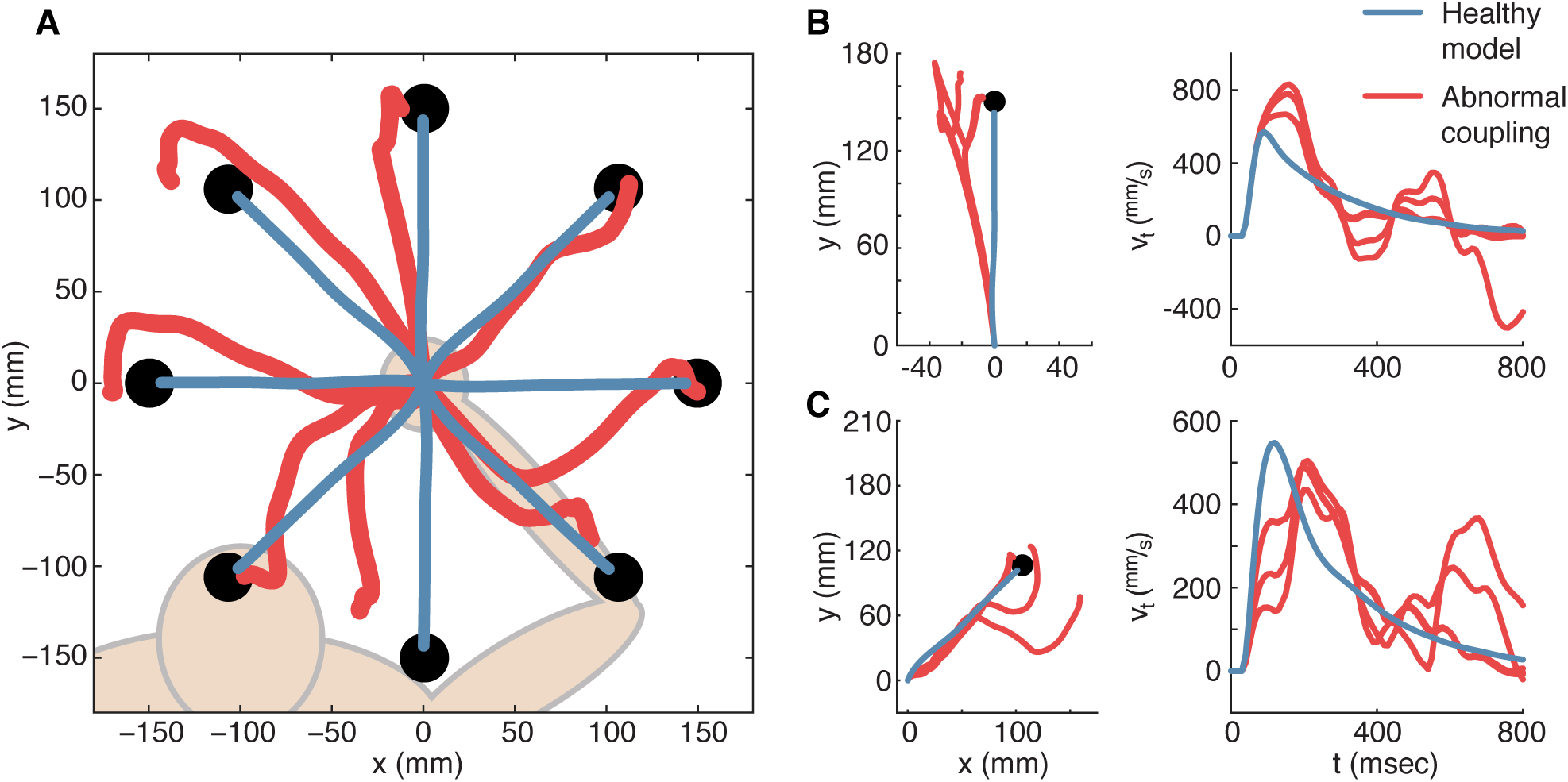
Effects of abnormal joint coupling on planar reaches. **Panel A** shows average hand trajectories followed by a healthy model (blue) and one with abnormal joint coupling (red) [3] in a center-out reaching task. While the healthy model follows nearly straight trajectories to each target, the abnormally coupled model follows much longer trajectories that sometimes curve away from or overshoot the targets. **Panel B** shows six separate reaches (one with healthy model, five with abnormal coupling) to a single target straight in front of the simulated subject (left) and the tangential velocity of each reach plotted against time (right). The model with abnormal coupling shows systematic perturbations to the left as the hand approaches the target. **Panel C** shows six separate reaches (one with healthy model, five with abnormal coupling) to a single target rotated 45 degrees (clockwise) from the one shown in panel B (left) and the tangential velocity of each reach plotted against time (right). The model with abnormal coupling sometimes shows clockwise perturbations midway through a reach before missing the target. The perturbed model consistently takes longer to perform each reach and requires corrective movements not needed by the healthy model.

While simulations are robust to increases in estimation noise (*σ_p_* = 0.02 to *σ_p_* = 2, Fig. 3, panel A), once the noise gets large enough (*σ_p_* = 8, Fig. 3), it displays a significant effect on movement trajectories, primarily at the end of a reach as the arm attempts to settle on the target. The resultant “wandering” behavior can be attributed to error compounded during successive estimations over the course of the reach. The resultant trajectories are qualitatively similar to reaches seen in experiments with hemiparetic patients [11]. The trajectories also resemble data collected from patients suffering from deafferentation, a condition that leaves them with almost non-existent sensory input [42], [43]. As seen by comparing panels A.1 and A.2 of Fig. 3, this difficulty settling on a target might vary with reach direction. The severity likely correlates with the relative contributions of each joint to the desired movement, as quantified by the arm’s effective mass/inertia over the course of the reach. The arm is most sensitive to noise in directions of lower effective mass [42]. Regardless, given that a small change in estimation noise can make reaches look healthy again, future research should consider designing rehabilitation protocols that help patients develop robustness to this noise.

**Fig. 3.**
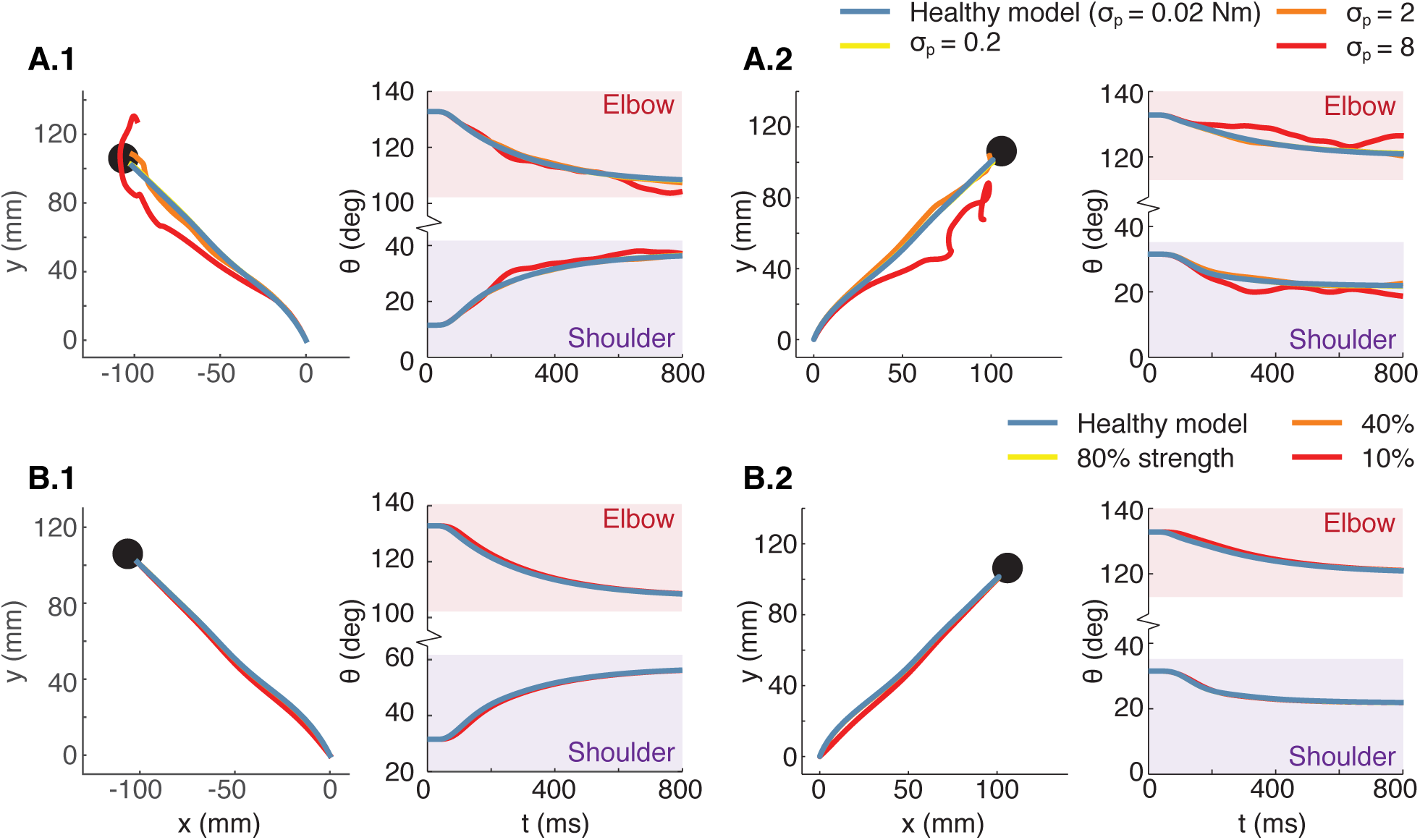
Sensitivity of reaching performance to estimation noise and muscular weakness. Post-stroke motor impairment frequently correlates with severity of the stroke. To determine the model’s sensitivity to increasing levels of sensorimotor impairment, we varied two model parameters. **Panel A** shows the results of increasing the internal model’s estimation noise—modeled as increased process noise in the internal model represented by the parameter *σ_p_* in our simulation framework—on planar reaches to two different targets in a center-out task. Yellow (*σ_p_* = 0.2*Nm*) and some orange (*σ_p_* = 2*Nm*) traces are hidden beneath blue traces (healthy model, *σ_p_* = 0.02*Nm*). As estimation noise increases, the internal prediction becomes less reliable (red), the reach deviates from the straight trajectory, and the hand requires more time to reach the target. However, reaching performance was not severely impaired by a two-order-of-magnitude change in this parameter from *σ_p_* = 0.02 (healthy) [29] up to *σ_p_* = 2. **Panel B** shows the effect of muscular weakness on two different planar reaches. Yellow (80% strength) and orange (40% strength) traces are hidden beneath the blue traces (healthy model). Weakness had no significant impact on reaching performance in this simulation.

The significance of muscular weakness for post-stroke motor impairment has been debated in the literature (e.g., [9], [44] versus [5], [10]). Our simulations display no motor impairment due to muscle weakness in a planar, gravity-supported reaching task. These findings agree with experimental studies in which stroke survivors who had undergone strength training therapy saw little improvement in motor capability [5], [10], [11]. However, muscular strength is likely to play a much larger role in motor performance in the presence of gravity or external loads (e.g., lifting objects).

Overall, our results indicate that reaching performance is highly robust to prediction error resulting from increased estimation noise, but is greatly impaired by unexpected joint coupling. More generally, the neuromuscular control system seems relatively insensitive to noise but very sensitive to unmodeled dynamics. This is likely thanks to the combination of Bayesian sensory integration [24] and optimization-based control that regularly re-plans motor commands [33]. Therefore, while many control models have focused on stochastic optimal control [45]–[47], a more difficult control problem appears to be the robust one: achieving adept motor performance in the face of poor internal models.

Though the framework presented here incorporates many biologically based parameters including mass/length/inertia properties, nerve conduction delays, and a near-optimal feedback controller thought to best approximate healthy neuromuscular control, some approximations were necessary. First, because nonlinear dynamics make the optimal control problem nonconvex [48], most simulations of upper-limb movements rely on linearized models for which optimal control is well-defined [49], [50]. Our model uses locally linear models that we assume to remain valid over the short (30-ms) time horizon used for MPC optimization. However, the human nervous system is likely able to plan movements over longer time horizons and account for dynamic nonlinearities. Second, while the unscented Kalman filter (UKF) [32] estimates arm state using the fully nonlinear dynamics, it requires the manual selection of several parameters. Third, the applied model perturbations are only approximations of hemiparetic sensorimotor impairments. For example, the abnormal coupling matrix was based on data from subjects performing maximum effort and may not scale linearly to smaller efforts. Despite such approximations, the unperturbed model behaves as expected, providing a tool to understand the map between sensorimotor deficits and motor performance. Model improvements and validation, both driven by data collected from stroke patients, will be the subject of future work.

While many of the sensorimotor deficits encountered by disabled stroke survivors are treatable, the benefits of applying a treatment cannot be foreseen due to the unknown causal relationship between sensorimotor deficits and motor impairments. A computational, parameterized simulation such as the one presented here allows for the selection of subject-specific parameters and sensorimotor deficits. In this way, it has the potential to simulate the effects of specific medical interventions so that only the most effective treatments are implemented on a patient. Additionally, the framework could allow for the optimal design of subject-specific assistive devices. While this has been done in healthy subjects [22], [23], technical limitations have prevented this type of simulated human-in-the-loop design of assistive devices for sensorimotor-impaired populations.

## List of Variables & Parameters

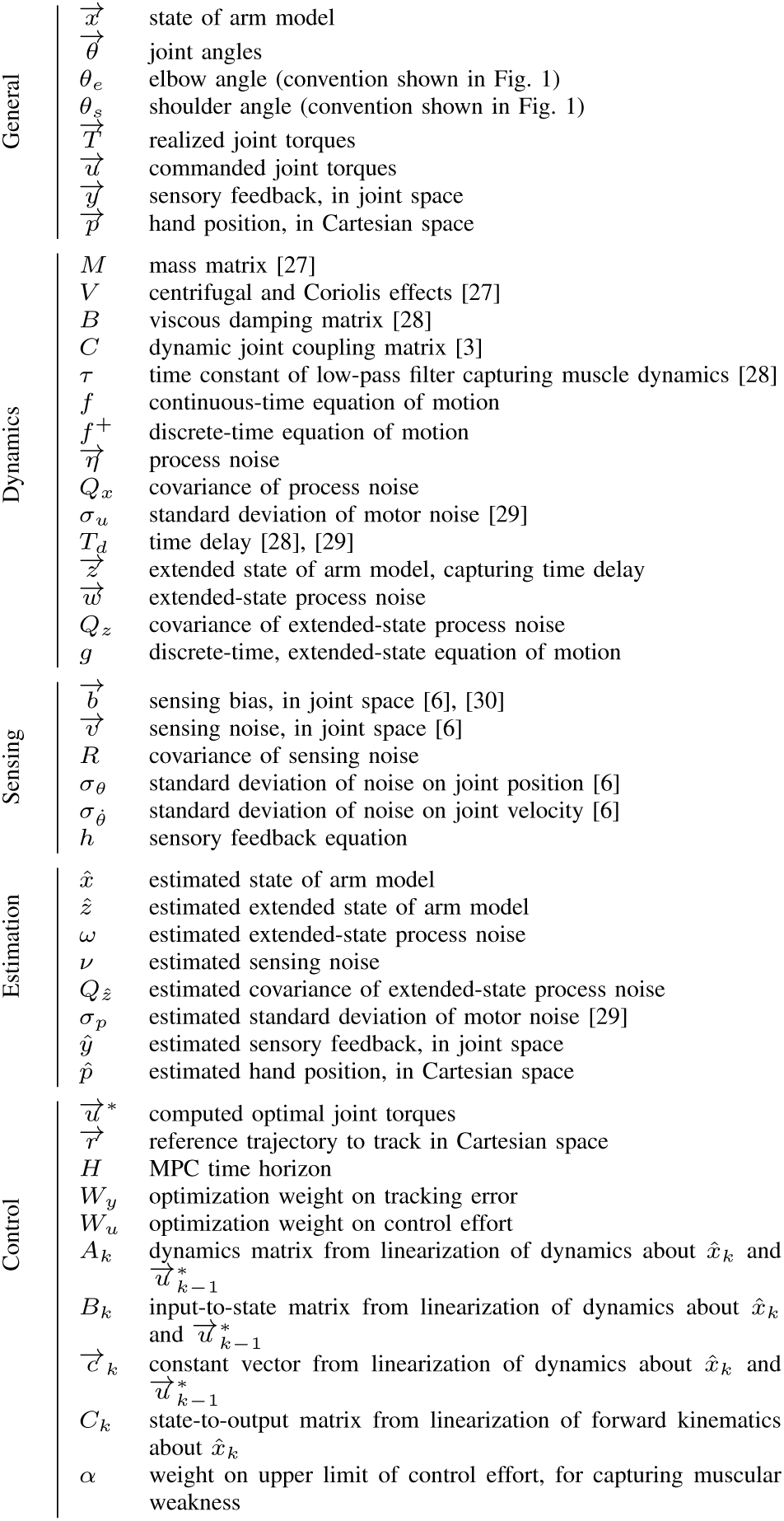

## REFERENCES

[1] D. W. Franklin, L. P. J. Selen, S. Franklin, and D. M. Wolpert, “Selection and control of limb posture for stability,” Proceedings of the Annual International Conference of the IEEE Engineering in Medicine and Biology Society, EMBS, pp. 5626–5629, 2013.

[2] J. J. Guilbert, “The world health report 2002 - reducing risks, promoting healthy life.” Education for Health, vol. 16, no. 2, p. 230, 2003.

[3] J. P. A. Dewald, P. S. Pope, J. D. Given, T. S. Buchanan, and W. Z. Rymer, “Abnormal muscle coactivation patterns during isometric torque generation at the elbow and shoulder in hemiparetic subjects,” Brain, vol. 118, no. 2, pp. 495–510, 1995.

[4] R. F. Beer, J. D. Given, and J. P. a. Dewald, “Task-dependent weakness at the elbow in patients with hemiparesis,” Archives of Physical Medicine and Rehabilitation, vol. 80, no. 7, pp. 766–772, 1999.

[5] J. P. Dewald and R. F. Beer, “Abnormal joint torque patterns in the paretic upper limb of subjects with hemiparesis.” Muscle & Nerve, vol. 24, no. 2, pp. 273–83, 2001.

[6] I. Cusmano, I. Sterpi, A. Mazzone, S. Ramat, C. Delconte, F. Pisano, and R. Colombo, “Evaluation of Upper Limb Sense of Position in Healthy Individuals and Patients after Stroke,” Journal of Healthcare Engineering, vol. 5, no. 2, pp. 145–162, 2014.

[7] D. Bourbonnais and S. V. Noven, “Weakness in patients with hemiparesis,” American Journal of Occupational Therapy, vol. 43, no. 5, pp. 313–319, 1989.

[8] C. L. Watkins, M. J. Leathley, J. M. Gregson, A. P. Moore, T. L. Smith, and A. K. Sharma, “Prevalence of spasticity post stroke,” Clinical Rehabilitation, vol. 16, no. 5, pp. 515–522, 2002.

[9] J. M. Wagner, C. E. Lang, S. A. Sahrmann, Q. Hu, A. J. Bastian, D. F. Edwards, and A. W. Dromerick, “Relationships between sensorimotor impairments and reaching deficits in acute hemiparesis.” Neurorehabilitation and Neural Repair, vol. 20, no. 3, pp. 406–416, 2006.

[10] K. M. Zackowski, A. W. Dromerick, S. A. Sahrmann, W. T. Thach, and A. J. Bastian, “How do strength, sensation, spasticity and joint individuation relate to the reaching deficits of people with chronic hemiparesis?” Brain, vol. 127, no. 5, pp. 1035–1046, 2004.

[11] R. F. Beer, M. D. Ellis, B. G. Holubar, and J. P. A. Dewald, “Impact of gravity loading on post-stroke reaching and its relationship to weakness,” Muscle and Nerve, vol. 36, no. 2, pp. 242–250, 2007.

[12] S. A. Chvatal and L. H. Ting, “Common muscle synergies for balance and walking.” Frontiers in Computational Neuroscience, vol. 7, no. May, p. 48, 2013.

[13] H. Müller and D. Sternad, “Decomposition of variability in the execution of goal-oriented tasks: three components of skill improvement.” Journal of Experimental Psychology: Human Perception and Performance, vol. 30, no. 1, pp. 212–233, 2004.

[14] K. M. Steele, M. M. van der Krogt, M. H. Schwartz, and S. L. Delp, “How much muscle strength is required to walk in a crouch gait?” Journal of Biomechanics, vol. 45, no. 15, pp. 2564–2569, 2012.

[15] S. L. Delp, F. C. Anderson, A. S. Arnold, P. Loan, A. Habib, C. T. John, E. Guendelman, and D. G. Thelen, “OpenSim: Open-source software to create and analyze dynamic simulations of movement,” IEEE Transactions on Biomedical Engineering, vol. 54, no. 11, pp. 1940–1950, 2007.

[16] H. Hultborn and J. B. Nielsen, “Spinal control of locomotion–from cat to man,” Acta Physiologica, vol. 189, no. 2, pp. 111–121, 2007.

[17] A. E. Kerdok, A. A. Biewener, T. A. McMahon, P. G. Weyand, and H. M. Herr, “Energetics and mechanics of human running on surfaces of different stiffnesses,” Journal of Applied Physiology, vol. 92, no. 2, pp. 469–478, 2002.

[18] E. Todorov and M. I. Jordan, “Optimal feedback control as a theory of motor coordination.” Nature Neuroscience, vol. 5, no. 11, pp. 1226–1235, 2002.

[19] E. Todorov, “Optimality principles in sensorimotor control,” Nature Neuroscience, vol. 7, no. 9, pp. 907–915, 2004.

[20] S. H. Scott, “Optimal feedback control and the neural basis of volitional motor control,” Nature Reviews Neuroscience, vol. 5, no. 7, pp. 534–546, 2004.

[21] F. C. Anderson and M. G. Pandy, “Dynamic optimization of human walking.” Journal of Biomechanical Engineering, vol. 123, no. 5, pp. 381–390, 2001.

[22] C. Ong, J. Hicks, and S. Delp, “Simulation-Based Design for Wearable Robotic Systems: An Optimization Framework for Enhancing a Standing Long Jump,” IEEE Transactions on Biomedical Engineering, vol. 63, no. 5, pp. 894–903, 2015.

[23] T. K. Uchida, A. Seth, S. Pouya, C. L. Dembia, J. L. Hicks, and S. L. Delp, “Simulating ideal assistive devices to reduce the metabolic cost of running,” PloS ONE, vol. 11, no. 9, p. e0163417, 2016.

[24] K. P. Körding and D. M. Wolpert, “Bayesian decision theory in sensorimotor control,” Trends in Cognitive Sciences, vol. 10, no. 7, pp. 319–326, 2006.

[25] J. Gordon, M. F. Ghilardi, S. E. Cooper, and C. Ghez, “Accuracy of planar reaching movements,” Experimental Brain Research, vol. 99, no. 1, pp. 112–130, 1994.

[26] J. W. Krakauer, M.-F. Ghilardi, and C. Ghez, “Independent learning of internal models for kinematic and dynamic control of reaching,” Nature Neuroscience, vol. 2, no. 11, pp. 1026–1031, 1999.

[27] D. A. Winter and H. J. Yack, “EMG profiles during normal human walking: stride-to-stride and inter-subject variability.” Electroencephalography and Clinical Neurophysiology, vol. 67, no. 5, pp. 402–411, 1987.

[28] F. Crevecoeur and S. H. Scott, “Priors Engaged in Long-Latency Responses to Mechanical Perturbations Suggest a Rapid Update in State Estimation,” PLoS Computational Biology, vol. 9, no. 8, p. e1003177, 2013.

[29] J. Izawa and R. Shadmehr, “On-Line Processing of Uncertain Information in Visuomotor Control,” Journal of Neuroscience, vol. 28, no. 44, pp. 11 360–11 368, 2008.

[30] C. T. Fuentes and A. J. Bastian, “Where is your arm? Variations in proprioception across space and tasks,” Journal of Neurophysiology, vol. 103, no. 1, pp. 164–171, 2010.

[31] R. Shadmehr and J. W. Krakauer, “A computational neuroanatomy for motor control,” Experimental Brain Research, vol. 185, no. 3, pp. 359–381, 2008.

[32] E. A. Wan and R. Van Der Merwe, “The unscented kalman filter for nonlinear estimation,” in Adaptive Systems for Signal Processing, Communications, and Control Symposium, 2000, pp. 153–158.

[33] M. Dimitriou, D. M. Wolpert, and D. W. Franklin, “The temporal evolution of feedback gains rapidly update to task demands,” Journal of Neuroscience, vol. 33, no. 26, pp. 10 898–10 909, 2013.

[34] E. K. Chadwick, D. Blana, R. F. Kirsch, and A. J. van den Bogert, “Real-Time Simulation of Three-Dimensional Shoulder Girdle and Arm Dynamics,” IEEE Transactions on Biomedical Engineering, vol. 141, no. 4, pp. 520–529, 2014.

[35] J. H. M. van Dijk, “Simulation of human arm movements controlled by peripheral feedback,” Biological Cybernetics, vol. 29, no. 3, pp. 175–186, 1978.

[36] R. D. Crowninshield and R. a. Brand, “A physiologically based criterion of muscle force prediction in locomotion.” Journal of Biomechanics, vol. 14, no. 11, pp. 793–801, 1981.

[37] M. Herceg, M. Kvasnica, C. Jones, and M. Morari, “Multi-Parametric Toolbox 3.0,” Proceedings of the European Control Conference, pp. 502–510, 2013.

[38] J. Roh, W. Z. Rymer, and R. F. Beer, “Evidence for altered upper extremity muscle synergies in chronic stroke survivors with mild and moderate impairment,” Frontiers in Human Neuroscience, vol. 9, no. 6, pp. 1–14, 2015.

[39] A. M. Coderre, A. A. Zeid, S. P. Dukelow, M. J. Demmer, K. D. Moore, M. J. Demers, H. Bretzke, T. M. Herter, J. I. Glasgow, K. E. Norman et al., “Assessment of upper-limb sensorimotor function of subacute stroke patients using visually guided reaching,” Neurorehabilitation and Neural Repair, vol. 24, no. 6, pp. 528–541, 2010.

[40] R. F. Beer, J. P. Dewald, and W. Z. Rymer, “Deficits in the coordination of multijoint arm movements in patients with hemiparesis: Evidence for disturbed control of limb dynamics.” Experimental Brain Research, vol. 131, no. 3, pp. 305–319, 2000.

[41] H. Jørgensen, H. Nakayama, H. Raaschou, and T. S. Olsen, “Stroke. Neurologic and functional recovery the Copenhagen Stroke Study.” Physical Medicine and Rehabilitation Clinics of North America, vol. 10, no. 4, pp. 887–906, 1999.

[42] C. Ghez, J. Gordon, M. F. Ghilardi, C. N. Christakos, and S. E. Cooper, “Roles of proprioceptive input in the programming of arm trajectories,” in Cold Spring Harbor Symposia on Quantitative Biology, vol. 55, 1990, pp. 837–847.

[43] C. Ghez, J. Gordon, M. F. Ghilardi, and R. L. Sainburg, “Contributions of vision and proprioception to accuracy in limb movements.” in The Cognitive Neurosciences, 1995, pp. 549–564.

[44] L. Ada, S. Dorsch, and C. G. Canning, “Strengthening interventions increase strength and improve activity after stroke: a systematic review,” Australian Journal of Physiotherapy, vol. 52, no. 4, pp. 241–248, 2006.

[45] D. W. Franklin and D. M. Wolpert, “Computational mechanisms of sensorimotor control,” Neuron, vol. 72, no. 3, pp. 425–442, 2011.

[46] E. Todorov, “Stochastic optimal control and estimation methods adapted to the noise characteristics of the sensorimotor system,” Neural Computation, vol. 17, no. 5, pp. 1084–1108, 2005.

[47] C. M. Harris and D. M. Wolpert, “Signal-dependent noise determines motor planning.” Nature, vol. 394, no. 6695, pp. 780–784, 1998.

[48] S. Boyd and L. Vandenberghe, Convex optimization. Cambridge University Press, 2004.

[49] E. Todorov and W. Li, “A generalized iterative LQG method for locally-optimal feedback control of constrained nonlinear stochastic systems,” Proceedings of the American Control Conference, pp. 300–306, 2005.

[50] W. Li and E. Todorov, “Iterative linear quadratic regulator design for nonlinear biologoical movement systems,” 1st International Conference on Informatics in Control, Automation and Robotics, pp. 1–8, 2004.

